# Whole genome sequencing of *Plasmodium falciparum* from dried blood spots using selective whole genome amplification

**DOI:** 10.1101/067546

**Authors:** Samuel O. Oyola, Cristina V. Ariani, William L. Hamilton, Mihir Kekre, Lucas N. Amenga-Etego, Anita Ghansah, Gavin G. Rutledge, Seth Redmond, Magnus Manske, Dushyanth Jyothi, Chris G. Jacob, Thomas D. Otto, Kirk Rockett, Chris I. Newbold, Matthew Berriman, Dominic P. Kwiatkowski

## Abstract

Translating genomic technologies into healthcare applications for the malaria parasite *Plasmodium falciparum* has been limited by the technical and logistical difficulties of obtaining high quality clinical samples from the field. Sampling by dried blood spot (DBS) finger-pricks can be performed safely and efficiently with minimal resource and storage requirements compared with venous blood (VB). Here, we evaluate the use of selective whole genome amplification (sWGA) to sequence the *P. falciparum* genome from clinical DBS samples, and compare the results to current methods using leucodepleted VB. Parasite DNA with high (> 95%) human DNA contamination was selectively amplified by Phi29 polymerase using short oligonucleotide probes of 8-12 mers as primers. These primers were selected on the basis of their differential frequency of binding the desired (*P. falciparum* DNA) and contaminating (human) genomes. Using sWGA method, we sequenced clinical samples from 156 malaria patients, including 120 paired samples for head-to-head comparison of DBS and leucodepleted VB. Greater than 18-fold enrichment of *P. falciparum* DNA was achieved from DBS extracts. The parasitaemia threshold to achieve >5x coverage for 50% of the genome was 0.03% (40 parasites per 200 white blood cells). Over 99% SNP concordance between VB and DBS samples was achieved after excluding missing calls. The sWGA methods described here provide a reliable and scalable way of generating *P. falciparum* genome sequence data from DBS samples. Our data indicate that it will be possible to get good quality sequence data on most if not all drug resistance loci from the majority of symptomatic malaria patients. This technique overcomes a major limiting factor in *P. falciparum* genome sequencing from field samples, and paves the way for large-scale epidemiological applications.

## Introduction

The last decade has seen rapid advances in whole genome sequencing technologies helping to track disease outbreaks and the spread of drug resistance genes[1–5]. Clinical and public health applications for *Plasmodium falciparum* sequencing rely on obtaining sequenceable material from samples collected in the field, often in resource-limited conditions. To date, the practical difficulties in sample collection, storage and transportation impose significant barriers to the use of genomic approaches for malaria surveillance.

The most practical and convenient method for sampling clinical malaria parasites is through small blood volumes obtained from capillary blood using finger or heel-pricks[6, 7]. These small blood samples – about 50 μl in volume - are blotted on filter papers for efficient transportation and storage without requiring refrigeration; this is especially applicable to resource-deprived regions where the disease is endemic. Despite the convenience and ease of sampling, DNA extracted from dried blood spot (DBS) filter papers often has low parasite DNA yield and an overwhelming host DNA contamination, which poses serious limitations in downstream genetic analyses[8]. These technical bottlenecks have prevented analysis of large numbers of pathogen samples collected by DBS at whole genome resolution, including archived clinical specimens, using current high throughput sequencing technologies. Currently, whole blood from malaria patients used for *P. falciparum* sequencing is obtained through venous blood (VB) draws. This requires skilled phlebotomists or clinicians with appropriate training. Once collected, VB samples are processed by filtering out leukocytes using cellulose columns[9] and require refrigerated storage followed by centrifugation and blood pellet freezing or DNA extraction. These requirements limit the scope for sample collection in remote regions where healthcare infrastructure is already under strain. The cellulose filtration process, although very effective in parasite enrichment, requires large volumes of blood (>2 ml)[10]. Such volumes can be difficult to obtain, especially from young children, who may already be anaemic as a result of *P. falciparum* infection[11] and who bear the heaviest disease burden globally.

To overcome the challenges of low sample quality and quantity, and to allow timely genetic analysis of clinical samples collected directly from patients without culture adaptation, we have used an approach that selectively amplifies parasite DNA from low blood volume clinical samples. The selective whole genome amplification (sWGA) strategy, originally described by Leichty and Brisson[12], uses computationally selected short oligonucleotide probes of 8-12 mers as primers that preferentially bind to the target genome, and this approach has been successfully applied to *Laverania* parasites including *P. falciparum* [13, 14]. The purpose of the present study was to undertake a detailed evaluation of sWGA approaches for sequencing the *P. falciparum* genome from dried blood spots.

## Material and Methods

### Primer design and selection

In order to design probes that preferentially bind to the *P. falciparum* genome, we used a published PERL script[12] to select up to 100 (8-12 mer) primers with a predicted specified melting temperature (≤ 30°C). We compared the frequency of these primer sequences in the desired (D) *Plasmodium falciparum* 3D7 genome against the contaminating (C) human genome (Figure 1A). The top 50 primers with the highest desired/contaminating (D/C) ratios were selected for further analysis. From these 50, primers with more than three complementary nucleotides at 3’ and 5’ ends were removed to prevent formation of hairpin structures. To prevent primer-primer dimerization, primer pairs with more than three complementary nucleotides at their ends were also removed. A final 28 primers that passed the above QC were ordered from Integrated DNA Technologies (Coralville, IA) as standard desalting purification with a single modification of phosphorothiorate bond between the last two 3’nucleotides to prevent primer degradation by the Phi29 polymerase exonuclease activity. Individual primers were reconstituted in Tris HCl (pH 8.0) buffer and pooled into three sets (probes) following the D/C ranking described above: the first set consisted of the first 10 primers (Probe_10), the second set consisted of the first 20 primers (Probe_20), and the third set consisted of all 28 primers (Probe_28). We evaluated each set to determine which should be taken forward for further assessment.

**Figure 1:**
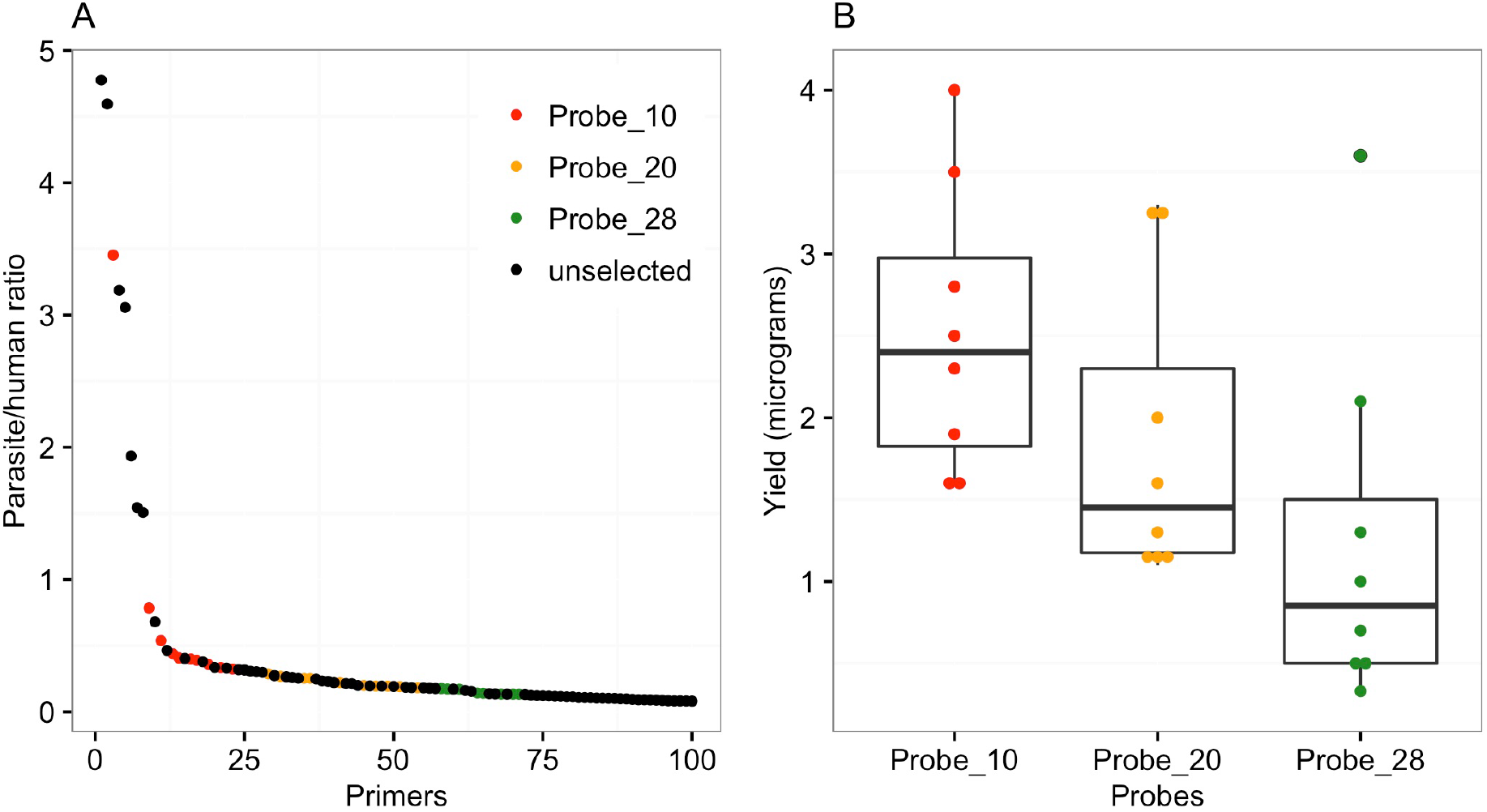
Primer selection analysis. A) The frequency of each primer as ranked by the frequency of occurrence. The y-axis represents the calculated ratio of the frequency in the parasite (desired) genome against frequency in the human (contaminating) genome. The x-axis shows the order of ranking by the frequency of occurrence ratio. B) DNA yield obtained following the selective whole genome amplification with different pools of probe sets. Probe_10 represent a pool of the first top 10 ranking primers. Probe_20 is a cumulative mixture consisting of all the Probe_10 primers plus the next 10 probes in that order. Probe_28 is a cumulative mixture of all the first 28 primers (Probe_10, Probe_20 and the next 8 probes in the order of frequency ranking).

### Mock samples to test the efficacy of sWGA

To test whether our selected primers would successfully amplify the parasite genome, we prepared mock clinical samples by mixing culture-infected red blood cells (infected with *P. falciparum* strain 3D7) with uninfected human whole blood to obtain simulated parasitaemias ranging from 0.0001% to 1%. *P. falciparum* strain 3D7 parasites for the mock samples were cultured in human O+ erythrocytes with heat-inactivated 10% pooled human serum, as described in [15]. All parasitaemia calculations were based on the estimation of approximately 4 million red blood cells per microliter of whole blood. Mock samples (N = 8) were used to extract DNA without leukodepletion. In addition, 6 other mock DNA samples were manually reconstituted by mixing *P. falciparum* genomic DNA with host (human) genomic DNA to obtain parasite/host DNA mixtures of the ratio 1: 24.

### Whole genomes from dried blood spots

To test the efficacy of sWGA to generate reliable genomic data from DBS, we compared WGS datasets obtained from standard leukodepleted-genomic DNA (gDNA) against sWGA-DBS samples. The gDNA samples were derived from two to three ml of venous blood with CF-11 or MN filtration to remove leukocytes[9, 10] prior to DNA extraction. The DBS samples were collected by spotting 50 ul whole blood obtained by finger pricking. In total, we analysed samples from 156 patients that were positive for *P. falciparum* clinical malaria based on rapid diagnostic test (RDT) with CareStart™ Malaria kit (Access Bio Inc, USA). Eighty-four DBS were collected from the Kassena-Nankana Districts of Upper East Ghana, of which 48 had matching VB pairs; and 72 DBS were collected from Noguchi Memorial Hospital in Accra, Ghana, all of which had matching VB samples.

### DNA extraction and quantification

Two to three ml of VB samples (mock blood or field) were used to extract DNA, using QiAamp DNA blood midi kit (Qiagen) following the kit manufacturer’s instructions. For DBS, DNA was extracted using QIAamp DNA Investigator Kit (Qiagen, Valencia, California, United States). Approximately 1.5 cm (0.6 inch) diameter DBS circles from each filter paper were cut out into small pieces of 3 mm diameter using a single-hole paper punch. Punched pieces from each sample were placed into 2 ml micro-centrifuge tubes from which DNA was extracted following the manufacturer’s instructions except for the reagent volumes and incubation times, which were doubled to accommodate the increased amount of DBS used per sample. An average of 116 ng (standard deviation, sd, 116.7) of DNA was obtained from the DBS extracts out of which at least 5 ng was used as template for sWGA amplification reaction.

### Selective whole genome amplification (sWGA)

The sWGA reaction was performed in 0.2 ml PCR-tubes or plates. The reaction (50 μl total volume) containing at least 5 ng of template DNA, 1x BSA (New England Biolabs), 1 mM dNTPs (New England Biolabs), 2.5 μM of each amplification primer, 1x Phi29 reaction buffer (New England Biolabs), and 30 units of Phi29 polymerase (New England Biolabs), was placed in a PCR machine (MJ thermal Cycler, Bio-Rad) programmed to run a “stepdown” protocol consisting of 35°C for 5 min, 34°C for 10 min, 33°C for 15 min, 32°C for 20 min, 31°C for 30 min, 30°C for 16 hrs then heating at 65°C for 15 min to inactivate the enzymes prior to cooling to 4°C. Once the product was amplified, it was quantified using Qubit® dsDNA high sensitivity (Thermo Fisher Scientific) to determine whether there was enough material for sequencing – minimum required is 500 ng of product. Standard whole genome amplified (WGA) products of the test samples were also sequenced as control to determine the extent of enrichment[16].

### Library preparation of amplified samples and short read high throughput sequencing

sWGA products (≥ 500 ng total DNA) were cleaned using Agencourt Ampure XP beads (Beckman Coulter) following manufacturer’s instructions. Briefly, 1.8 volumes of beads per 1 volume of sample were mixed and incubated for 5 min at room temperature. After incubation, the tube containing bead/DNA mixture was placed on a magnetic rack to capture the DNA-bound beads while the unbound solution was discarded. Beads were washed twice with 200 μl of 80% ethanol and the bound DNA eluted with 60 μl of EB buffer. Cleaned amplified DNA products (∼ 05 – 1 μg DNA) were used to prepare a PCR-free Illumina library using the NEBNext DNA sample preparation kit (New England Biolabs) for high throughput sequencing. DNA libraries were sequenced at the Wellcome Trust Sanger Institute using Illumina HiSeq 2500 instruments and Illumina V.3 chemistry. Paired-end sequencing was performed with 100-base reads and an 8-base index read. 12-multiplex sample libraries were loaded to target at least 20 million reads per sample.

### Data analysis

Sequence data obtained from each sample was subjected to standard Illumina QC procedures and 20 million reads per sample was subjected to detailed analysis for enrichment, quality, content, and coverage. Each dataset was analysed independently by mapping sequence reads to the 3D7 reference genome using BWA[17]. SAMtools[18] was used to generate coverage statistics from the BWA mapping output. For enrichment analysis we counted the number of reads mapping to either host, or *P. falciparum* reference sequences. For genotype and concordance analysis, we generated variation calls using SAMtools (V0.1.1.19) mpileup and bcftools (V0.1.17). We used a list of 1,241,840 (1.2 million) high-quality single-nucleotide polymorphism (SNP) positions, part of which had been published[19, 20] and then performed *in silico* genotyping of both the DBS (sWGA) and VB (gDNA) samples using mpileup to count alleles present in at least five reads. SNP call concordance analysis between matching DBS and VB samples was performed on sequenced data targeting key malaria drug resistance genes, such as *crt* (K76T involved in chloroquine resistance)[21], *dhfr* (N51I, involved in pyrimethamine resistance)[22], *dhps* (A581G, involved in sulfadoxine resistance)[23], *mdr1* (N86Y, involved in multiple drugs including mefloquine)[24], and *kelch13* (C580Y, involved in artemisinin resistance)[25].

## Results

### sWGA primer selection and amplification yield

We analysed our selected 28 primers individually (Figure 1A) to determine their expected binding sites and distribution pattern across the *P. falciparum* genome. Each 1 or 2kb block had at least one primer binding (Figure S1). We then pooled the selected 28 primers into three different sets (probes): Probe_10 (consisting of the first 10 primers), Probe_20 (consisting of the first 20 primers), and Probe_28 (a pool of all the 28 primers). In separate reactions, we used the three probes to amplify 5 ng of simulated mock samples (a mix of 3D7-infected red blood cells with uninfected human whole blood) to determine which set gives optimal genome amplification and coverage. The amplified products were cleaned and the DNA quantified using Quant-iT™ PicoGreen® dsDNA assay kit (Invitrogen) to determine the yield for each primer pool (Figure 1B). We observed different yields between the three primer pools: Probe_10 produced the highest average yield (2.5 ± 0.87 μg) followed by Probe_20 (1.85 ± 0.81 μg) and Probe_28 (1.2 ± 1.0 μg) (Figure 1B). We then performed whole genome sequencing of the amplified products and compared the quality of genome coverage (number of bases with at least 5x coverage) by each set (pool) and found no significant difference (Spearman’s correlation: Probe_10 and Probe_20, R^2^ = 0.97, *p* < 0.001; Probe_10 and Probe_28, R^2^ = 0.96, *p* < 0.001; Probe_20 and Probe_28, R^2^ = 0.97, *p* < 0.001). We therefore chose Probe_10 for all subsequent sWGA reactions based on amplification yield and cost.

### Coverage profile of sWGA samples

To perform a more in-depth analysis on Probe_10, we amplified and sequenced a mock sample containing a mixture of human (96%) and *P. falciparum* (4%) DNA as described in Methods. Using the Illumina short read sequence data obtained, we plotted the primer binding positions as well as the short-read sequence alignments against the reference genome using Circos software[26] for data visualisation. Figure 2 shows the probe binding sites (middle circle) as well as the sequence reads coverage profile (outermost circle) on all 14 chromosomes (inner circle) of the *P. falciparum* genome. Probe_10 successfully amplifies the majority of the parasite genome, but the variable subtelomeres are not adequately covered (Figure 2 and Figure S1). However, the coverage profile is uneven and does not correlate properly with the primer binding sites (Pearson’s correlation R^2^ = –0.007, *p* = 0.3). Previous analysis [19] revealed accessible and inaccessible regions in the *P. falciparum* genome. Inaccessible regions, mainly the telomeres, centromeres and sub-telomeres, are comprised of hypervariable and/or highly repetitive sequences that are difficult to assemble or map. The remaining parts (core genome) consist of mainly the coding sequences of relatively balanced-base composition, and are generally accessible in most genome analysis. In order to test whether sequences generated from sWGA samples would successfully cover the core genome, we plotted the coverage profile of *P. falciparum* strain 3D7 samples sequenced as gDNA (3 samples without amplification), WGA DNA (3 samples amplified using optimised whole amplification method[16]) or sWGA (3 samples consisting of a mixture of 4% parasite and 96% human DNA, amplified using selective whole genome amplification method). Figure 3 shows the coverage profile of the samples on chromosome 1, highlighting regions corresponding to the core genome. gDNA provided the most even and uniform coverage across the entire genome. Both WGA and sWGA samples produced relatively spiky and uneven coverage, with the sWGA producing higher coverage depths of uneven distribution.

**Figure 2:**
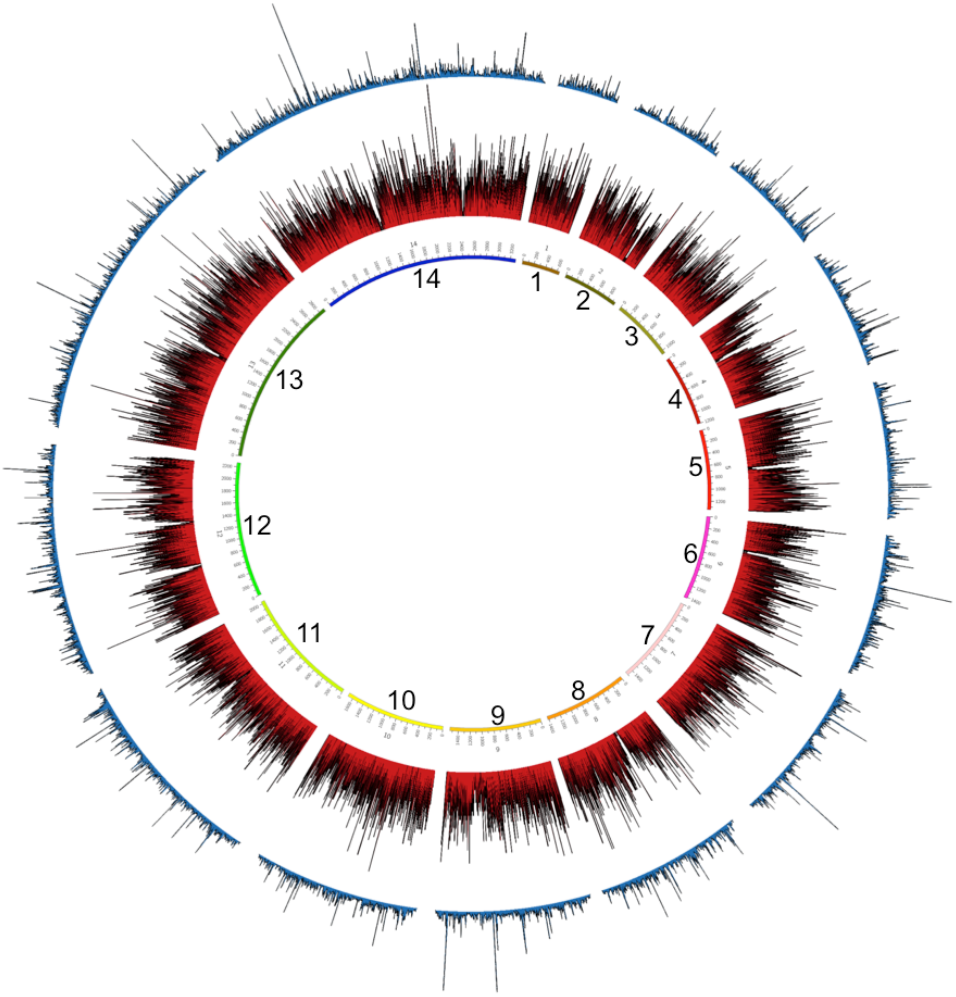
Circos plot analysis of Probe_10 primer pool and *P. falciparum* genome coverage. The three rings represent, from innermost to outermost, the 14 *P. falciparum* chromosomes and position in kb, the total number of primers binding in 1kb windows (red lines), and the average read depth in 1kb windows (blue lines). The figure was generated using the Circos software[26].

**Figure 3:**
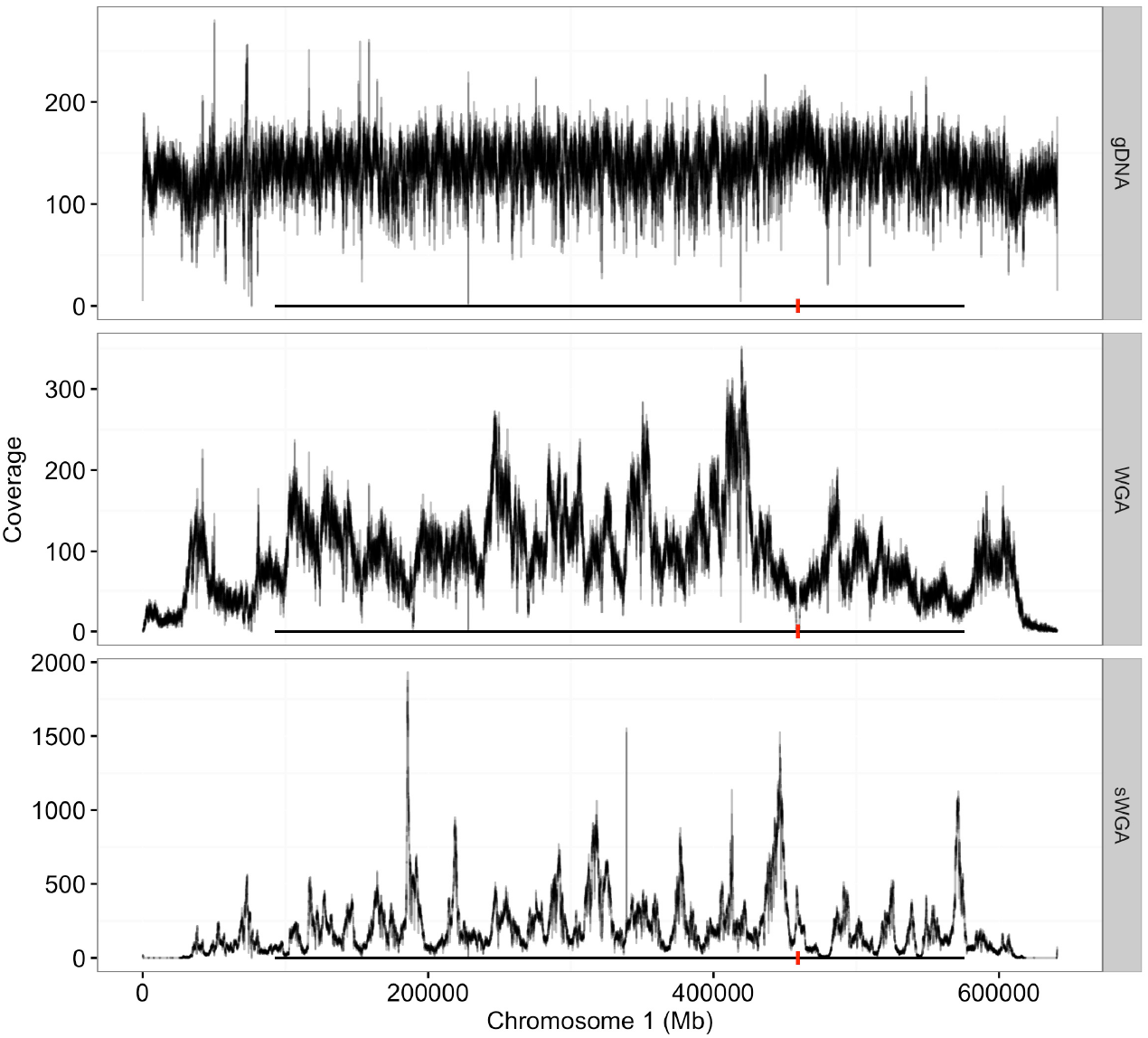
Core genome coverage profile. Coverage depth of chromosome 1 of unamplified (gDNA), whole genome amplified (WGA) and selective whole genome amplified (sWGA) DNA of *P. falciparum* strain 3D7. Black horizontal line shows positions corresponding to the core genome and red vertical line shows the centromere.

### sWGA enriches for *Plasmodium falciparum* sequence reads

We analysed the sequence data by comparing datasets from standard whole genome amplified (WGA) samples against their selectively amplified (sWGA) counterparts to determine the level of enrichment. An average of 73.2% (sd 4.4) of the reads in the sWGA-treated samples mapped to *P. falciparum*. More than 18-fold enrichment of parasite DNA was achieved, depending on the extent of host contamination in the original sample (Table 1; Figure 4). In contrast, data obtained from DBS extracts and amplified by standard WGA (no selective amplification) had less than 1% of reads mapping to *P. falciparum* and the rest (>99%) mapping to the host genome (Figure 4), demonstrating the efficacy of sWGA in selective amplification of parasite DNA.

**Figure 4:**
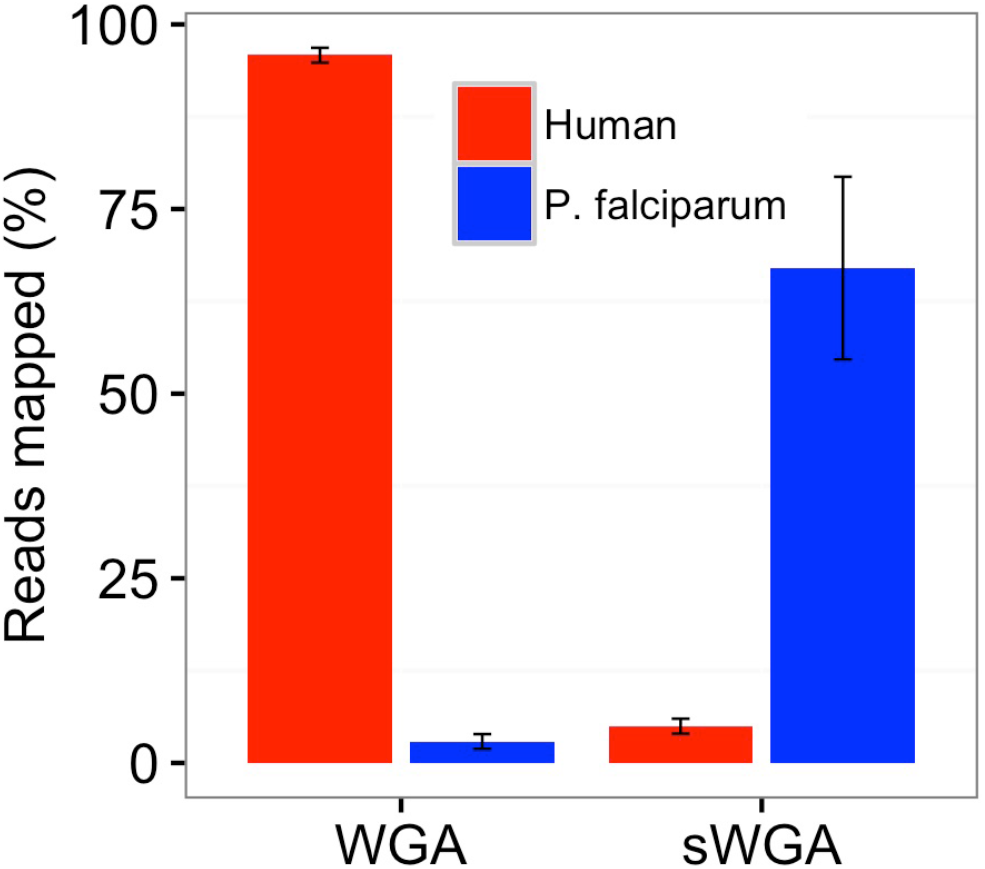
Selective whole genome amplification (sWGA) enrichment. Simulated clinical samples comprising 96% human DNA and 4% *P. falciparum* DNA (3D7) were amplified using either WGA or sWGA. Amplified samples were sequenced to determine the proportion of reads mapping to human or *P. falciparum* reference genomes.

**Table 1:** sWGA enrichment analysis. Mock and field samples were amplified by either WGA or sWGA before sequencing. Proportions of reads mapping to either human or *P. falciparum* genomes were used to determine the level of parasite DNA enrichment by sWGA treatment. 3D7 represent mock samples prepared by mixing P. falciparum and human genomic DNA in the ratio of 1:24. Field represent clinical genomic DNA samples extracted from dried blood spot filter papers.

| Reads mapping to: |  |  |  |  |
| --- | --- | --- | --- | --- |
| Sample | *P. falciparum* (%) | Human (%) | Others (%) | Fold Enrichment |
| WGA_3D7_1 | 3.30 | 96.20 | 0.50 | NA |
| WGA_field_1 | 2.90 | 95.83 | 1.27 | NA |
| sWGA_3D7_10 | 79.74 | 3.33 | 16.94 | 19.93 |
| sWGA_3D7_20 | 74.64 | 4.82 | 20.54 | 18.66 |
| sWGA_Field_10 | 73.50 | 4.45 | 22.04 | N/A |
| sWGA_Field_22 | 69.26 | 4.81 | 25.93 | N/A |
| sWGA_Field_28 | 68.84 | 5.55 | 25.61 | N/A |

### Parasitaemia and genome coverage threshold in mock samples

To investigate the sensitivity of the sWGA application, we analysed genome coverage threshold by sequence data generated from samples with different levels of parasitaemia. *In vitro* infected red blood cells were mixed with human whole blood to simulate different levels of clinical parasitaemia ranging from 1.0 to 0.0001%. For samples with parasitaemias of ≥ 0.005% (∼6.25 parasites per 200 white blood cells (WBC)), ≥ 70% of the core nuclear 3D7 genome was covered at depth of ≥ 5x reads. However, the coverage dropped sharply for samples with parasitaemia below 0.005% (Figure 5A; see Figure S2 for detailed coverage distribution). The same dataset was used to analyse coverage of known important drug resistant loci in the genome[2]. As shown in Figure 5B, a similar coverage profile was observed where all the 7 specified drug resistant loci were covered 100% at depths of ≥5x reads for samples with parasitaemia ≥ 0.005%.

**Figure 5:**
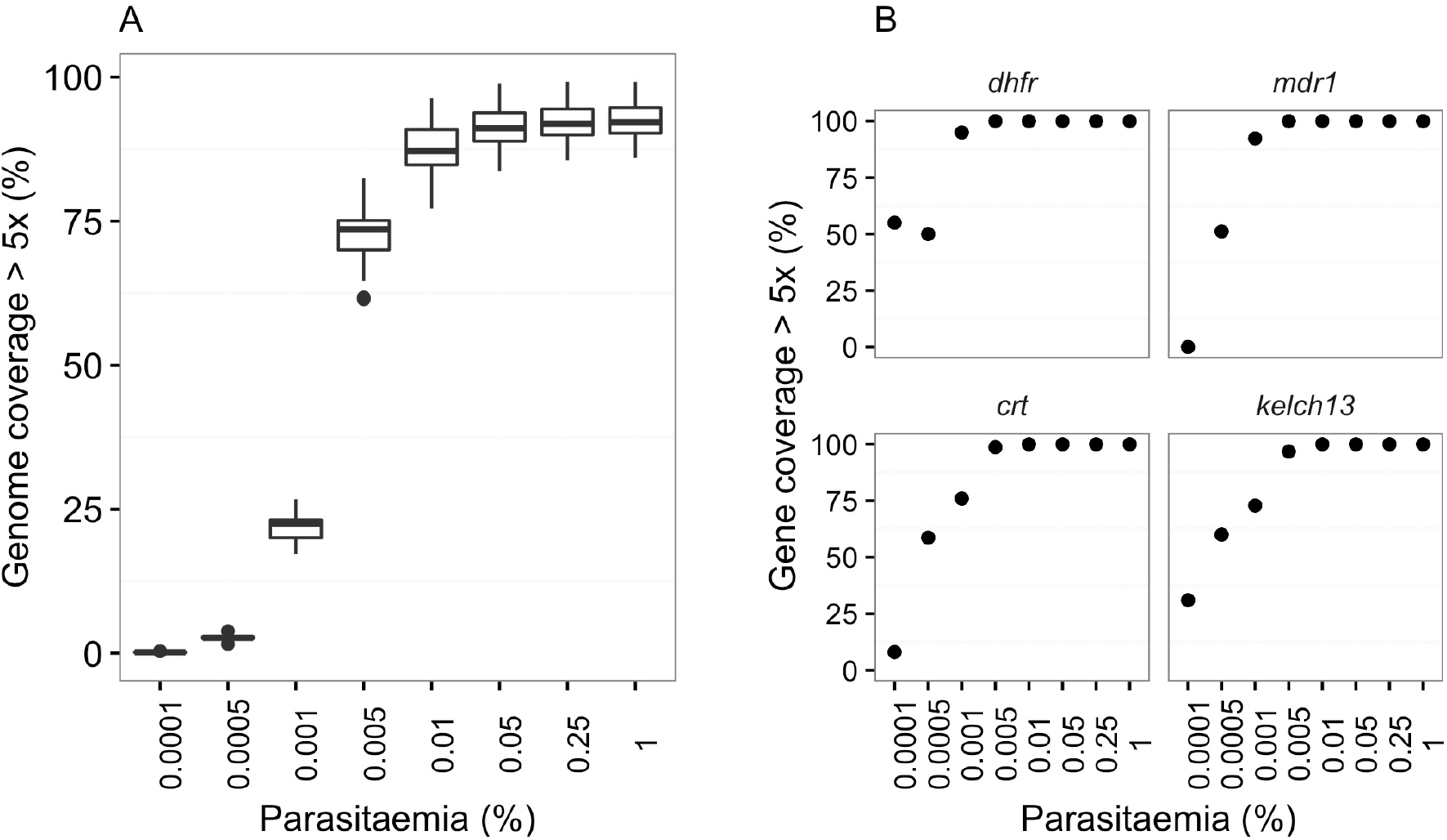
Assessing sWGA sensitivity and parasitaemia threshold. Clinical mock samples represent different levels of parasitaemia ranging from 0.0001% to 1%. Data from sWGA-processed samples were analysed to determine coverage of *P. falciparum* genome. A) Genome coverage by samples of different parasitaemia levels. B) Coverage of important drug resistance loci by mock samples of different levels of parasitaemia.

### sWGA allows whole genome sequencing directly from clinical dried blood spots

Having established sWGA efficacy in mock blood samples, we tested the method using DBS field isolates collected from two sites in Ghana, with parasitaemias ranging from 0.001% to 8.9% (1.25 to 11,125 parasites per 200 WBC or 40 to 356,000 parasites per μl of blood). We extracted DNA from 205 DBS samples (average yield 116 ng, sd 116.7), which were subsequently subjected to sWGA (average yield 1,399 ng, sd 502). From those, 156 (76%) passed the threshold of 500 ng for library preparation and were therefore whole genome sequenced.

We analysed 156 DBS samples and excluded those with less than 50% of the core genome covered at 5x reads or less (N = 25). On average only 2.3% (sd 2.3) of the core genome of the 131 DBS samples was not covered at all (Figure 6A), whereas 85% (sd 13) of the core genome was covered at 5x or more (Figure 6B). The median coverage of the core genome was 29x (Figure 6C).

**Figure 6:**
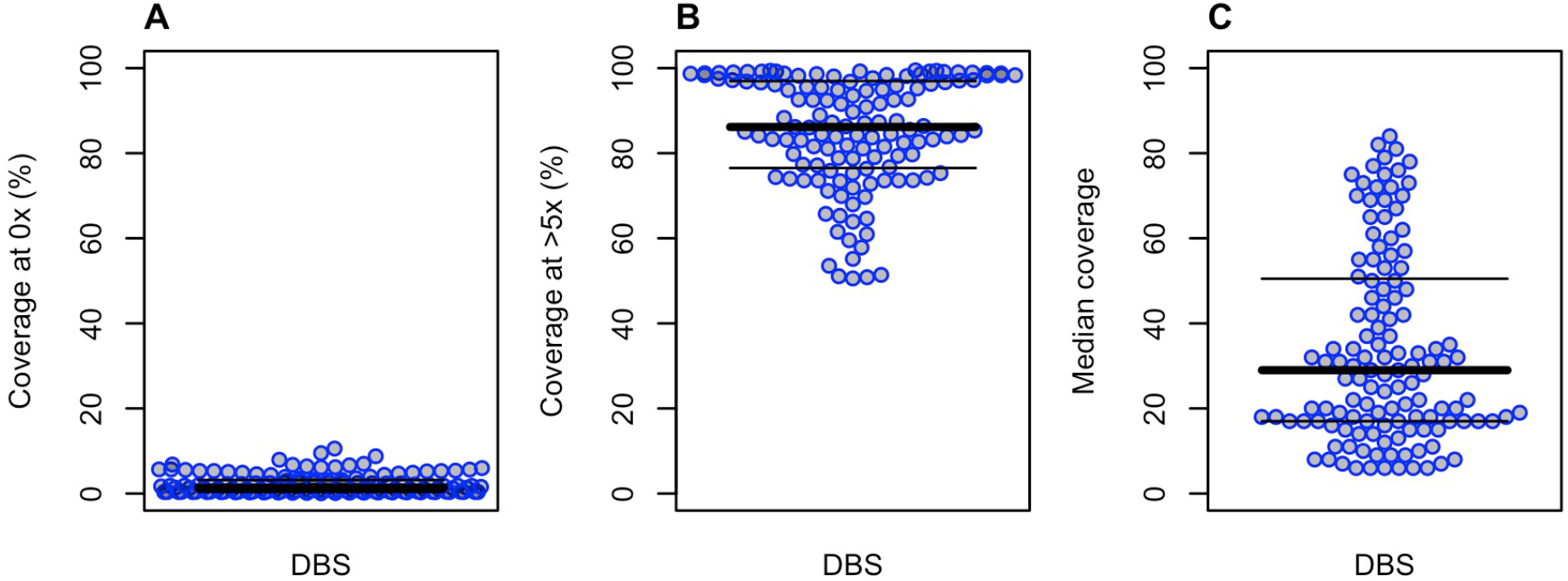
Core genome coverage of 156 dried-blood spot (DBS) clinical samples subjected to sWGA. A) Percentage of genome positions with no coverage. B) Percentage of genome positions with at least 5x coverage. C) Median genome coverage.

As expected, samples with higher parasitaemia (above 0.1%) produced sequence data with better coverage at depths of ≥5x, whereas samples with parasitaemia lower than 0.03% had many positions covered at depths less than 5x (Figure 7, *F* (_1,150_) = 135.5, *p* < 0.001). In our dataset all samples with parasitaemia lower than 0.03% (N = 25) had more than 50% of the genome covered at 5x or less. There was one exception; a sample with 0.001% parasitaemia had 51.4% of the core genome covered at 5x. The samples with low parasitaemia had a much larger proportion of missing bases in the core genome (Figure 7). We analysed the coverage of genes that are either responsible for, or associated with, antimalarial drug resistance (Figure S3), and observed a general tendency of better coverage in samples of higher parasitaemia (> 0.02%), while those with parasitaemia lower than 0.01% showed poor coverage across the genes.

**Figure 7:**
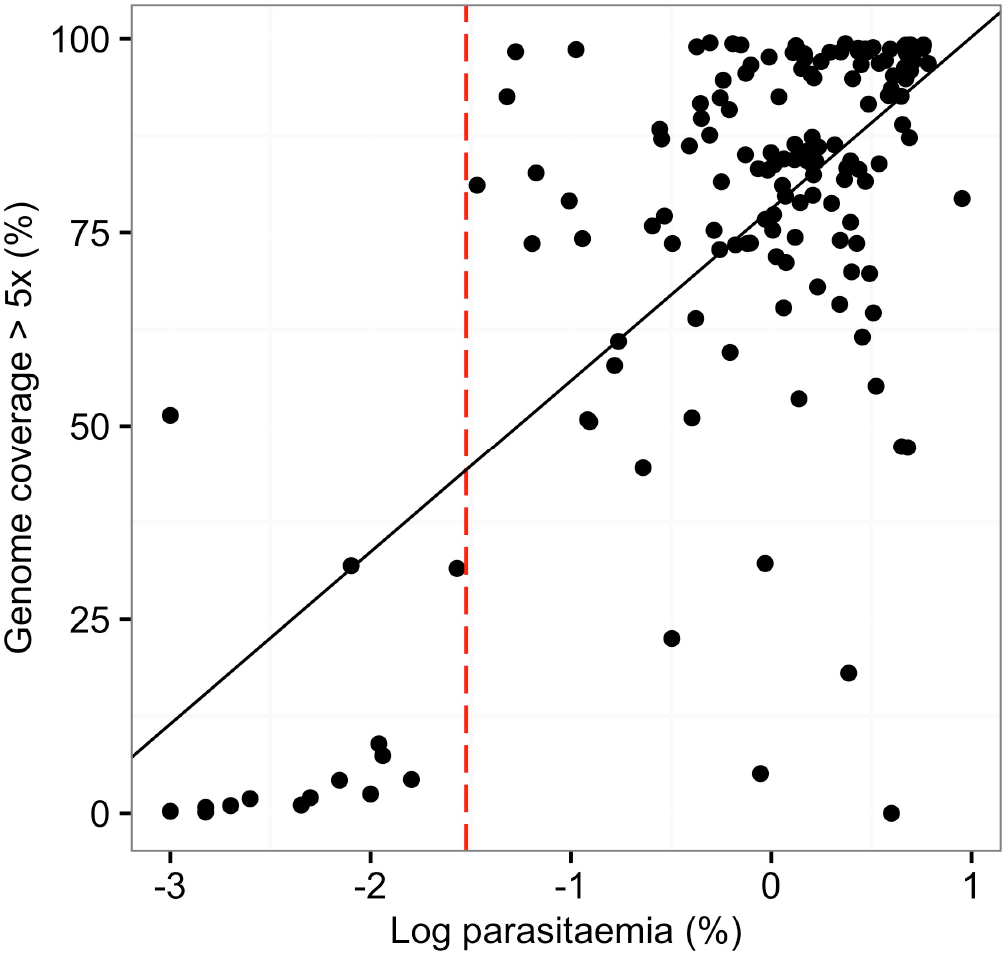
Field DBS samples with higher parasitaemia have a higher proportion of genome positions covered at depth greater than 5x. (*F* _(1-150)_ = 135.5; p = 0.001). The parasitaemia threshold to obtain at least 50% of the genome covered at 5x is 0.03% (dotted red line).

Taken together, our data establishes 0.03% parasitaemia (40 parasites per 200 WBC) as the minimum threshold on which sWGA technology is capable of generating quality sequence data with coverage suitable for most genetic analyses on DBS field samples (Figure 7, vertical dotted line marks the 0.03% parasitaemia threshold). We also established that at least 180 parasite genomes per sample is required for efficient sWGA processing.

### High concordance between dried blood spot samples and venous blood samples

In order to further evaluate sWGA efficiency and suitability for genetic studies from DBS samples we performed a concordance analysis using the set of 120 field samples with matched pairs of VB and DBS filter papers. Sequence data from both VB and DBS samples were analysed in parallel and genotyped against the ∼ 1.2 million high quality SNP positions previously identified in the *P. falciparum* genome[19]. Genotype calls from matching VB (gDNA) and DBS (sWGA) sample pairs were analysed. In the gold standard VB samples, a median of more than 98 % (N = 1,217,003) of SNPs were called in all the samples (Figure 8). Overall, 93% (N = 1,154,911) of SNPs were called in the DBS samples, with a slight reduction at lower parasitaemias (Figure 8). We then investigated the accuracy of the SNPs called by performing a concordance analysis between SNP calls made from VB and DBS samples. We excluded the samples that had less than 50% of the genome covered at 5x (N = 7) as well as all missing calls of the remaining DBS samples (total samples analysed N=113). There was high concordance between the SNPs called in both VB and DBS samples, with an average of more than 99.9% (out of 1,241,840 SNPs, sd 10.97%) of calls being concordant (either Ref/Ref or Alt/Alt; Table 2). Only 0.04% (out of 1,241,840 SNPs, sd 0.08%) of calls were discordant (Ref/Alt or Alt/Ref).

**Figure 8:**
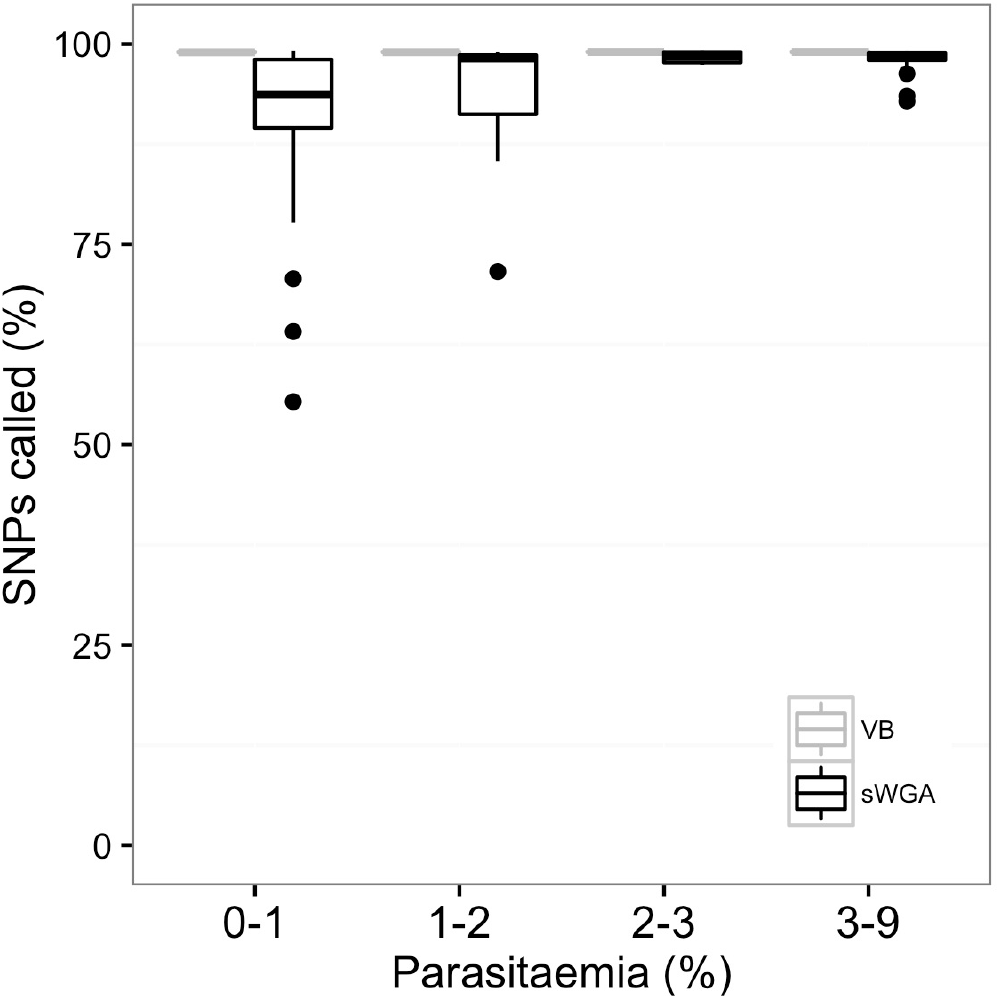
Single nucleotide polymorphism (SNP) calls from matching venous blood samples (VB) and dried blood spots samples (DBS). The percentage of SNPs called in DBS samples decreases as parasitaemia decreases.

**Table 2:** sWGA and gDNA SNP concordance analysis (N=113). Genotype calls across ∼1.2 million biallelic typable SNPs from matching venous blood (gDNA) and DBS (sWGA) sample pairs were analysed to obtain SNP concordance between the two sample processing methods. Ref, reference genotype call; Alt, alternative genotype call; Het, heterogeneous calls; Miss, missing calls.

|  | **Ref/Ref** | **Alt/Alt** | **Ref/Alt** | **Alt/Ref** |
| --- | --- | --- | --- | --- |
| Average | 1,130,936.8<br>(99.82 %) | 1,391.3<br>(0.12 %) | 288.4<br>(0.02%) | 283.4<br>(0.02%) |
| SD | 124,553.8<br>(10.9%) | 848.6<br>(0.07 %) | 477.8<br>(0.04%) | 498.3<br>(0.04 %) |
| Median | 1,179,088<br>(99.9 %) | 1,026<br>(0.08%) | 0 | 0 |

We further tested the accuracy of SNP calls from sWGA-generated data using allele frequency concordance metrics. Using the VCF files targeting the 1.2 million high quality-biallelic SNPs, we analysed the population-level allele frequencies from matching VB and DBS samples and found strong correlation of non-reference allele frequencies (NRAF) between VB (gDNA) and DBS (sWGA) samples (Figure 9; Pearson’s correlation *P* = 0.99, *p* < 0.001). For more detailed analysis, we looked at the allele frequencies of specific mutations for key malaria drug resistance genes - *dhfr, mdr1, crt, dhps* and *kelch13*. Once again, we found high concordance between VB and DBS samples (Figure 9, Table S1). In summary, after excluding lower quality samples with missing calls, we observed very high concordance in population genetic data between VB and DBS samples.

**Figure 9:**
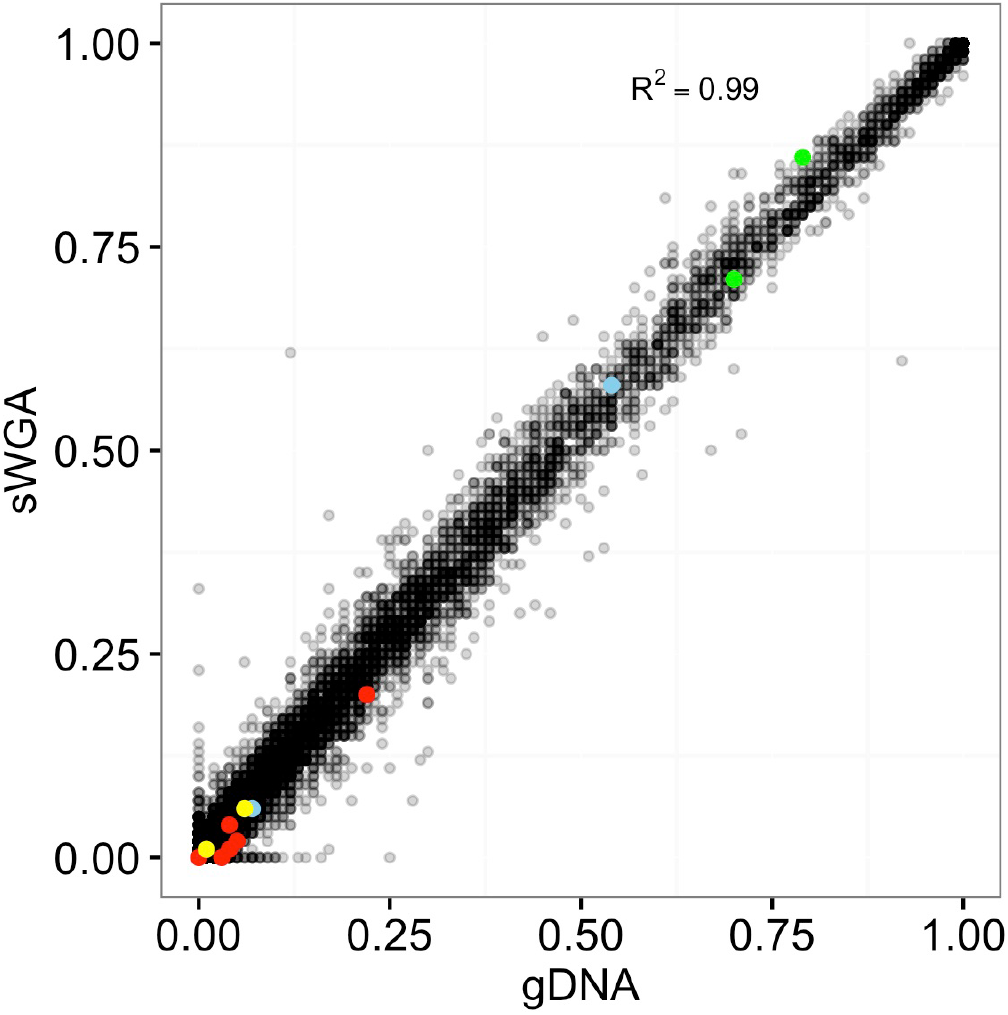
Population-level allele frequency of VB (gDNA) and DBS (sWGA) are strongly correlated. Using variant calls from the 1.2 million high quality-biallelic SNP positons, we compared non-reference allele frequencies (NRAF) from matching VB and DBS samples and obtained a strong correlation between the two sample sets. Coloured dots represent specific mutations of drug resistance genes. Green dots represent *dhfr* (N51I, C59R, S108N); blue dots represent *mdr1* (N86Y, Y184F); red dots represent *crt* (M74I, N75D, N75K, K76T, A220S, Q271E, I356T, R371I); yellow dots represent *dhps* (S436A, G437A, K540E, A581G, A613S).

## Discussion

Using sWGA to amplify parasite DNA from dried blood spots has immediate and important implications for public health. Here we have set out to comprehensively characterise the potential of the method. Collecting clinical malaria samples as DBS on filter paper is field-friendly and has several advantages for both patient and researcher over the venous blood (VB) draw methods currently used for parasite whole genome sequencing[27]. Finger-prick sampling requires less advanced training than VB draws, collects ∼50x less blood, and is more convenient for most patient groups. Unlike VB draws, DBS samples also do not require special facilities for transportation, refrigeration, and storage, since the blotted paper is stabilised by the membrane that preserves genetic integrity[27, 28]. VB sampling is thus relatively limited in geographic range, restricted to locations with well-established and resourced clinics. Sequencing from DBS samples would break this technical bottleneck, allowing significant expansion of sample collection to include very remote regions, increasing sampling density and coverage[27, 28].

We found that ∼110 ng of genomic DNA can be extracted from a 20 – 40 μl DBS, of which over 98% is host material. Isolating sequenceable parasite DNA from a DBS sample that is highly contaminated with host DNA has hindered applications of genetic tools in malaria research and control programmes. Previous studies have identified various methods to overcome the challenges of host DNA contamination in pathogen sequencing[8, 29]. However, most of these techniques require relatively large quantities of starting DNA material that is impossible to obtain from DBS samples. The approach we describe here provides a timely solution to this challenge, creating opportunities for both large-scale field isolate sequencing studies and analysing archived clinical samples that would otherwise be too contaminated and low yield for whole genome sequencing.

To thoroughly evaluate the quality and accuracy of sWGA sequence data for genetic analysis of clinical malaria samples, we sequenced and analysed 156 DBS collected from clinical malaria patients using sWGA. 120 of these had their corresponding VB counterparts collected simultaneously, allowing direct comparison between DBS (sWGA) and VB (gDNA) WGS data from an identical patient cohort[9, 10]. More than 75% of the *P. falciparum* genome was covered at ≥5x in 117 (97.5%) DBS samples for which parasitaemia was ≥0.03%. The sWGA-derived genome sequences show a less uniform coverage profile compared to data generated from unamplified genomic DNA (VB-derived, Figure 3). This is typical of whole genome amplified data[16]. However, the core genome was adequately covered at depths suitable for most downstream analysis including variant detection and SNP genotyping.

The high concordance of SNP calls and allele frequencies between the DBS and VB paired samples indicates that samples that were subjected to sWGA are suitable for population genetic studies. Significantly for the potential applicability of this technology to public health surveillance projects, important malaria drug resistance loci were successfully sequenced and showed very similar allele frequencies for both DBS and VB samples (Figure 9, Table S1).

## Conclusion

In summary, we have shown that processing DBS samples using sWGA method produces reliable sequence data, provided that: the sample has ≥ 180 *P. falciparum* genomes (parasitaemia threshold ∼0.03%, or ∼40 parasites per 200 WBC); the threshold for library preparation is met (≥500 ng of DNA post-sWGA); and the sequence data obtained covers at least 50% of the genome at a depth of 5x or more. Samples with much lower parasitaemias, for example collected from asymptomatic patients or during the low transmission season, will require further optimisation to improve sensitivity and coverage. Using sWGA technology, genomic data from larger sample sizes with geospatial resolution could provide useful information to public health bodies, for example through rapid detection of emerging patterns in parasite evolution in response to control initiatives such as antimalarial drugs.

## Declarations

### Ethical approval

The scientific merit and use of human subjects for this study was approved by the scientific review committee and ethical review boards of the Navrongo Health Research Centre of (FWA00000250) and the Noguchi Memorial Institute for Medical Research (056-12/13). Written informed consent was obtained from all adult subjects and from the parent or legal guardians of minors.

### Consent for publication

There are no case presentations that require disclosure of respondent’s confidential data/information in this study.

### Availability of data and materials

All data sets used in this study have been deposited in the European Nucleotide Archive read archive under accession number []

### Authors’ contributions

SOO, CVA, TDO, CIN, MB, and DPK conceived and coordinated the study; SOO, CVA, WLH, KR and MK performed the experiments; LNAE, AG, WLH and MK participated in sample collection and field experiments; SOO, CVA, WLH, GGR, MM, CJ, SR, CGJ and DJ participated in data analysis and interpretation of the results; SO, CVA, WLH and DPK participated in the drafting, editing and final preparation of manuscript. All authors read and approved the final manuscript.

## Acknowledgement

We are grateful to Roberto Amato for useful discussions on analysis. We thank the field workers who helped collect the clinical samples, working with the Navrongo Health Research Centre and Noguchi Memorial Institute for Medical Research/ University of Ghana.

## Supplementary figures and table

**Figure S1:** A plot of primers (probes) and their binding distribution on the *P. falciparum* genome. The topmost panel show cumulating binding positions and distribution profile of all the 28 primers. Black dots (1) show positions where the primer binds and Red (0) dots shows positions with no primer binding.

**Figure S2:** Coverage depths frequencies of samples with different parasitaemia levels

**Figure S3:** Coverage of genes associated with drug resistance. Colours reflect the percentage of genome covered, ranging from 5x (grey) to 30x or more (red).

**Table S1:** Non-reference allele frequencies (NRAF) of major drug resistance genes for VB (gDNA) and DBS (sWGA) samples. Gene name, chromosome number, position, mutation name, mutation type and the NRAF found in West Africa populations (MalariaGen https://www.malariagen.net/apps/pf/4.0/) are shown. Notably, the studied populations have high *dhfr* mutation frequencies and rare *crt* mutations. Presumably because the use of SP was widespread and is still being used (e.g. pregnancy prophylaxis), whereas enough time has passed since chloroquine was widely used [30–32].

